# Diverse ecological pressures govern microbial heterogeneity across dark ocean ecosystems

**DOI:** 10.64898/2026.11.22.767178

**Authors:** Katherine M. Klier, Marguerite V. Langwig, James C. Kosmopoulos, Rika E. Anderson, Karthik Anantharaman

## Abstract

The deep ocean regulates Earth’s biogeochemical cycles, yet accelerating warming, deoxygenation, and biodiversity loss threaten to fundamentally disrupt these processes. Despite microorganisms’ central role in global elemental cycling, deep-sea microbial communities remain poorly characterized, limiting our capacity to predict ecosystem responses to ongoing environmental perturbation. Although ocean circulation connects marine ecosystems across vast distances in the absence of physical barriers, secondary factors including depth, redox chemistry, and substrate availability create distinct microbial habitats. A key unresolved question is how these ecological pressures differentially shape microbial community structure beyond dispersal alone. Here, we examine the microbial ecology of two biogeochemically distinct yet spatially connected marine habitats at the East Pacific Rise (EPR) spreading center in the Pacific Ocean: deep-sea hydrothermal plumes and oxygen minimum zones. Using serial filtration coupled with community metagenomics, we reconstruct >700 metagenome-assembled genomes (MAGs) and perform comprehensive taxonomic, metabolic, and abundance profiling across both systems. We reveal pronounced community differentiation driven by redox chemistry, as well as underappreciated ecological pressures that exert strong and previously overlooked structuring influence on these communities. Our findings establish that deep-ocean microbial ecology is governed by a complex hierarchy of environmental forces extending well beyond physical dispersal, with direct implications for forecasting microbial responses to accelerating ocean deoxygenation and climate change.

## INTRODUCTION

The ocean is essential to all life on Earth, acting as a major carbon sink (1), supplying atmospheric oxygen (2), and supporting global economies (3). Underpinning nearly all oceanic processes are microorganisms, including bacteria, archaea, and viruses, that produce oxygen, fix carbon, serve as the base of marine food webs, and drive biogeochemical cycles (4). Growing recognition of the importance and versatility of marine microorganisms has spurred advances in ecology, biotechnology, and environmental monitoring (5–7). Despite this progress, vast regions of the deep ocean remain unexplored, leaving major gaps in our understanding of the diversity, function, and resilience of marine microbial communities (8–10).

While often referred to as a singular ecosystem, the ocean is not a homogenous basin, but a mixture of ecosystems defined by differences in factors such as depth, temperature, and chemistry. Despite the absence of physical barriers, these factors create distinct environmental gradients that persist into the dark ocean depths, exerting unique ecological pressures on the microorganisms within them. For instance, oxygen minimum zones (OMZs) are regions within the water column where dissolved oxygen (DO) concentrations are at their lowest, typically below 20μM (11,12). Here, some microorganisms are exposed to high influxes of organic matter from the photic zone which they can use for respiration, thus contributing to oxygen depletion (13). Others, in turn, must adapt to the resulting hypoxic/anoxic conditions through anaerobic metabolisms (12,14). On the seafloor, hydrothermal vents represent another extreme habitat where geothermally heated, chemically enriched fluids are discharged from the ocean floor into the surrounding seawater. Hydrothermal fluids not only expose microorganisms to extreme temperature gradients and highly reducing environments locally but they can also disperse over vast distances through hydrothermal plumes, influencing broader ocean ecology and geochemistry (15–17).

Despite their differences, these two environments are not wholly isolated. Currents and thermohaline circulation drive constant movement and mixing between waters of different depths and geographic locations (18,19). As a result, not only water itself, but also chemicals, substrates, and organisms are transported throughout the ocean (20–22). Hydrothermal plumes have also been documented to travel hundreds of kilometers from their source (23), and the fluid nature of the ocean means that microorganisms could, in principle, move freely between vent systems and OMZs. Consistent with this, studies have identified taxonomic overlap between microbes inhabiting hydrothermal vents and open ocean environments such as OMZs, reflecting connectivity (17,24). Others, in turn, have demonstrated clear depth-dependent shifts in marine microbial metabolism and composition (25,26), reflecting ecological stratification. Nevertheless, a direct OMZ-to-plume comparison has yet to be conducted despite the ubiquity of these ecosystems across global oceans. As both OMZs and hydrothermal vents are subject to ongoing change or expansion (2,27), it is important to determine whether shifts in environmental parameters will drive local microbial community divergence, or whether the fluid nature of the ocean will act as a buffer against drastic changes.

In this study, we directly compare the microbial communities from an OMZ and hydrothermal plume environment that are in close geographic proximity to determine if the microbial communities significantly diverge and, where differences are noted, infer the ecological pressures driving these changes. To achieve these goals, we sampled microorganisms from an OMZ and hydrothermal plume environment at the East Pacific Rise (EPR) spreading center, a tectonically active region in the Pacific Ocean (Figure 1a) (28). Using a serial filtration approach combined with metagenomics, we analyzed microbial taxonomy, distribution, abundance, metabolism, and phylogeny across these two environments. EPR represents one of the most extensively studied hydrothermal systems on Earth with over four decades of geological, geochemical, and ecological research and has recently served as the site of numerous microbial discoveries and insights (28–31). Despite this extensive research, metagenomics-based characterizations at EPR remain remarkably scarce. Therefore, in addition to providing ecological insights, our study provides the most comprehensive microbial genomic catalog of this system to date, transforming a landmark geological site into a reference point for deep-sea microbial ecology.

**Figure 1.**
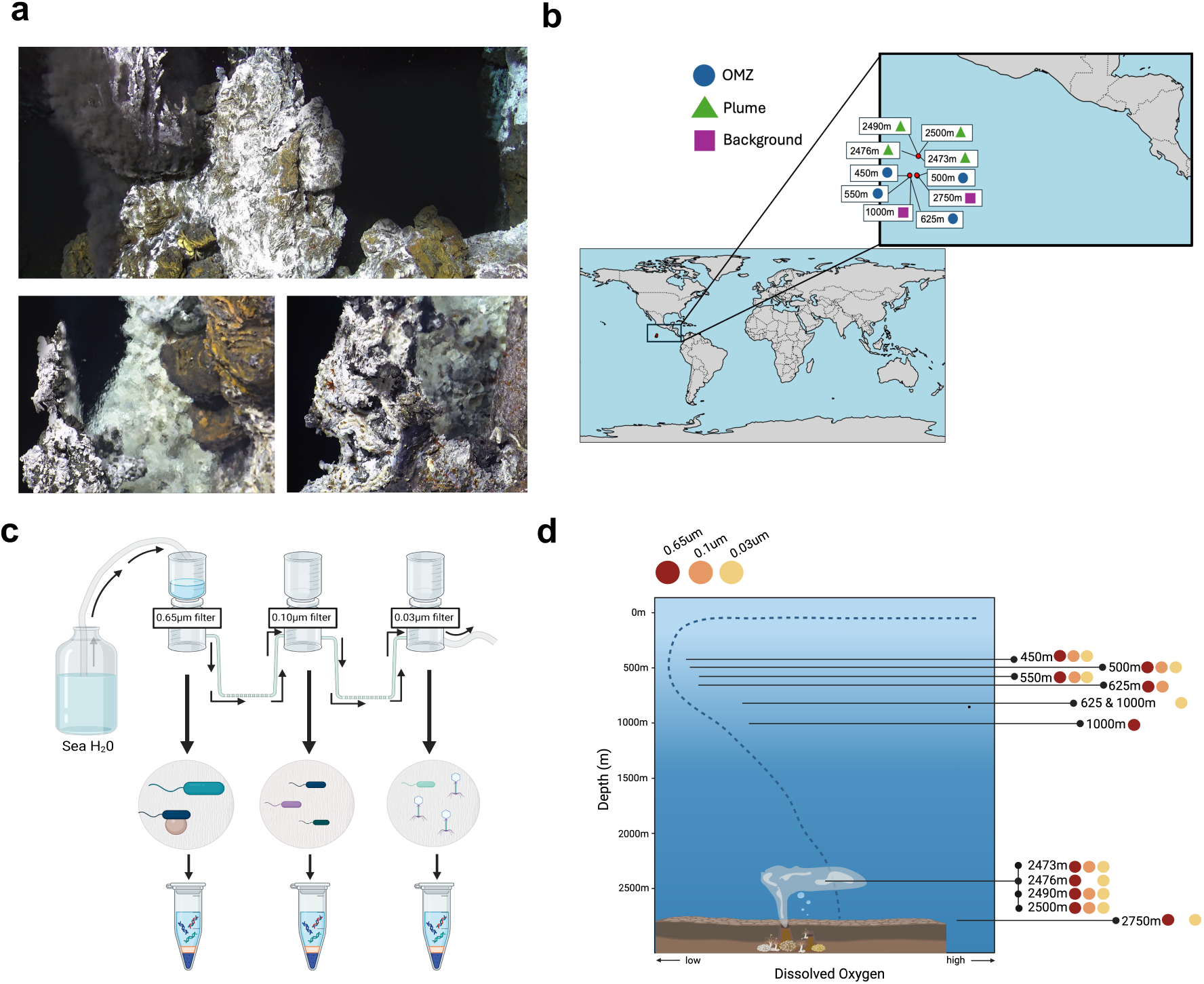
Schematic of microbial sampling methodology at EPR. a) Images of active venting sites at EPR taken at or near the time of sampling during cruise AT42-06 in December 2018. b) Geographic locations of sampling sites at EPR along with the specific depth that was sampled at each site. c) Schematic of the serial filtration approach used to sample and generate metagenomic assemblies at EPR. d) Representative illustration of a typical dissolved oxygen profile at EPR along with depths sampled and filter-size metagenomic assemblies that were successfully sequenced at each depth.

Our results reveal that while broad taxonomic overlap occurs between the two environments, community differentiation exists and becomes increasingly apparent at finer taxonomic resolution suggesting that environmental selective pressures act to “fine-tune” microbial communities rather than restructure them wholesale. We further demonstrate that the taxonomy or abundance of individual lineages can be a misleading proxy for environmentally driven differentiation, and that other community parameters, including metabolic strategy and cell size, diverge significantly between environments. By contextualizing differences within the conditions of each habitat, we identify a suite of potentially underappreciated forces, including thermodynamic constraints and viral predation, that together generate substantial heterogeneity in marine microbial communities.

## RESULTS

### Recovery of diverse microbial communities from the OMZ and hydrothermal plumes

Four depths at EPR were sampled within the OMZ (450m, 500m, 550m, and 625m), four within hydrothermal plumes (2473m, 2476m, 2490m, and 2500m), and two in background waters outside of both features (1000m and 2750m) (Figure 1b; Supplemental Table 1; Supplemental Figure 1). To distinguish larger or particle-associated microorganisms from smaller free-living cells and viruses, we employed a serial filtration approach using 0.65µm, 0.1µm, and 0.03µm filters in sequence. Independent metagenomic assemblies were generated for every filter fraction at each depth (Figure 1c). Metagenomic assemblies were successfully generated from the 0.65µm filter fraction for all 10 sampled depths, from the 0.1µm fraction for 7 sampled depths, and from the 0.03µm fraction for 9 sampled depths (Figure 1d; Supplemental Table 2), yielding assemblies from 26 distinct samples. Medium-to-high-quality MAGs (≥50% completion, ≤10% contamination (32)) were recovered from at least two filter fractions at all depths (Supplemental Table 2). Some 0.03µm assemblies yielded no medium-to-high-quality MAGs, likely due biomass loss following sequential filtrations.

Bacterial MAGs from the 0.65µm and 0.1µm fractions showed broadly similar phylum-level compositions, dominated by Pseudomonadota, Chloroflexota, and Marinisomatota (Figure 2ab; Supplemental Table 3). Distinctions in bacterial taxonomic composition between the larger two filter sizes at the phylum level included greater representation of MAGs from phylum Patescibacteria in the 0.1µm fraction, whereas MAGs representing some phyla, such as Gemmatimonadota and Verrucomicrobiota, were unique to the 0.65µm fraction. At finer taxonomic resolutions, additional differences emerged (Supplemental Figure 2ab; Supplemental Table 3). For example, within the phylum Pseudomonadota, orders *Pseudomonadales* and *Arenicellales* were the most highly represented order from the 0.65µm metagenomes, whereas MAGs from these two orders were absent from 0.1µm metagenomes.

**Figure 2.**
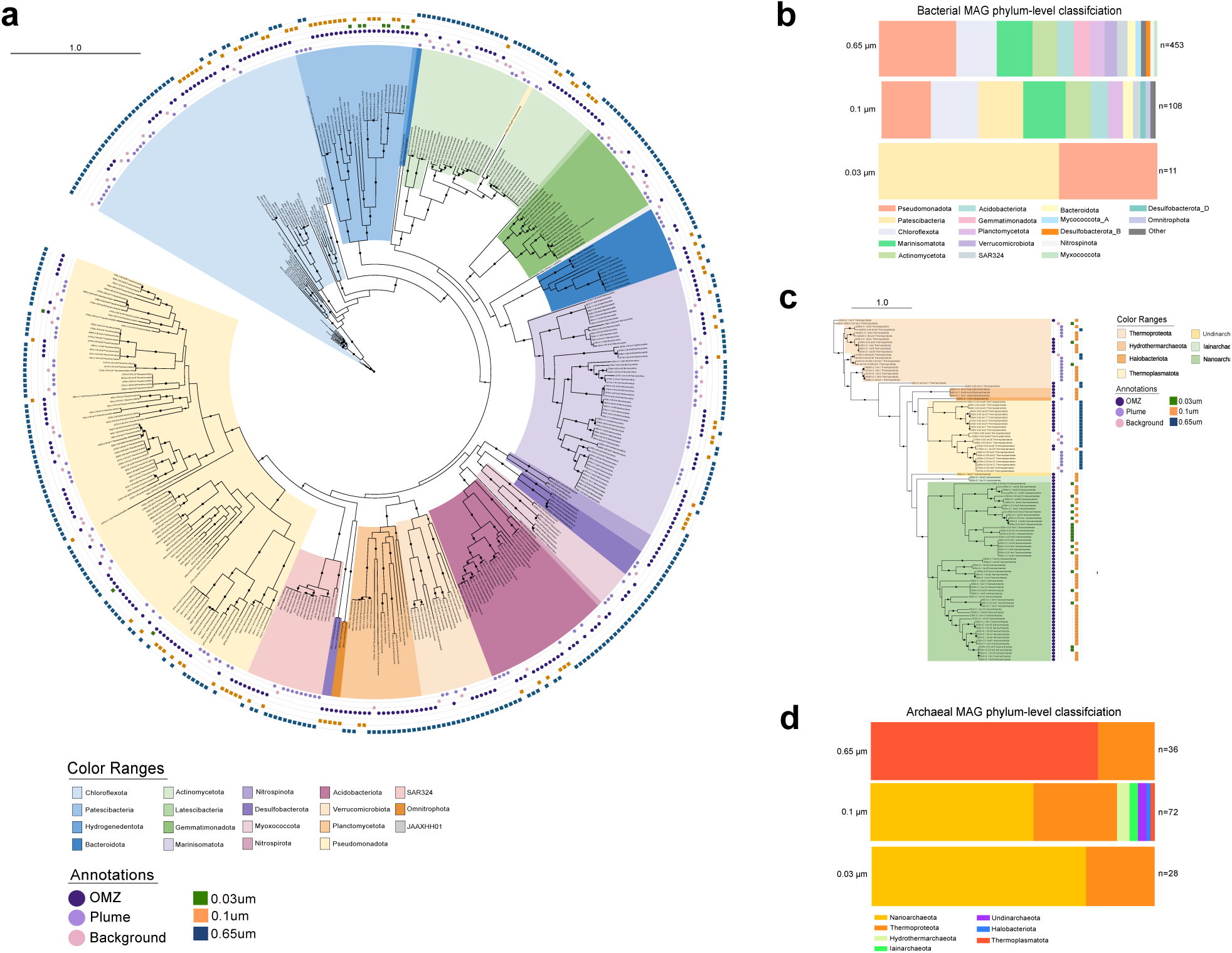
Taxonomy and distribution of bacterial and archaeal MAGs recovered from EPR across different filter sizes. a) Phylogenetic tree based on concatenated ribosomal subunit protein sequences from bacterial MAGs generated in this study that contained sufficient ribosomal protein sequencing data. The different color ranges indicate distinct phyla-level clusters. Each MAG is annotated with the filter size (squares) and the environment (circles) from which it originated. Black circles on tree branches indicate bootstrap values >90. b) Break down of bacterial MAG taxonomy by filter size across all depths, based on number of MAGs recovered from that phylum. Phyla represented by <1% of recovered MAGs from that filter size were grouped into an “other” category. Full classification information for all MAGs is available (Supplemental Table 3). The number of MAGs in each category is indicated to the right of the respective bar charts. c-d) The same as in a-b but for archaeal MAGs.

Archaeal MAGs showed greater phylum-level differentiation between fractions (Figure 2cd; Supplemental Table 3). The 0.65µm fraction was dominated by Thermoplasmatota, whereas the 0.1 µm fraction contained a more diverse assemblage, including Thermoproteota, Nanoarchaeota, Iainarchaeota, and Undinarchaeota.

Although the 0.03µm fraction was expected to contain primarily viral DNA, it yielded 11 bacterial and 28 archaeal MAGs (Figure 2a-d; Supplemental Table 2-3). While some recovery may reflect cell breakage during filtration, estimated MAG completion rates were comparable across the filter fractions but estimated genome size decreased (Supplemental Figure 3ab; Supplemental Table 4), suggesting that small intact cells can on occasion pass through filters smaller than the 0.22µm and 0.1µm pore sizes frequently utilized in environmental microbiome analyses (33–35). Dereplication at 95% average nucleotide identity (ANI) of MAGs across the three filter-sizes within respective sampling depths decreased MAG counts by, at most, 11% (Supplemental Figure 3c), supporting low microbial species redundancy across filter sizes.

Viral scaffolds were identified from all assemblies and comprised a larger proportion of the metagenomes from the 0.03µm filter fractions than the 0.1µm and 0.65µm fractions (Supplemental Figure 3d; Supplemental Table 5), indicating that the 0.03µm fraction was particularly enriched in viral sequences. Overall, these results demonstrate that sequential filtration recovered diverse microbial MAGs across size fractions while enriching viral sequences in the 0.03 µm fraction.

### The OMZ and plume support distinctive microbial communities

To determine if environmental selection pressures were present and evident based on taxonomic composition, we compared the taxonomic composition of microbiomes from the OMZ and hydrothermal plumes (Figure 3a-e). While background waters were sampled at depths of 1000m and 2750m, a 0.1µm fraction metagenome was not successfully recovered from either background sample. To avoid artificially inflating dissimilarity due to missing data, background waters were excluded from comparisons involving consolidation of data from various filter sizes.

**Figure 3.**
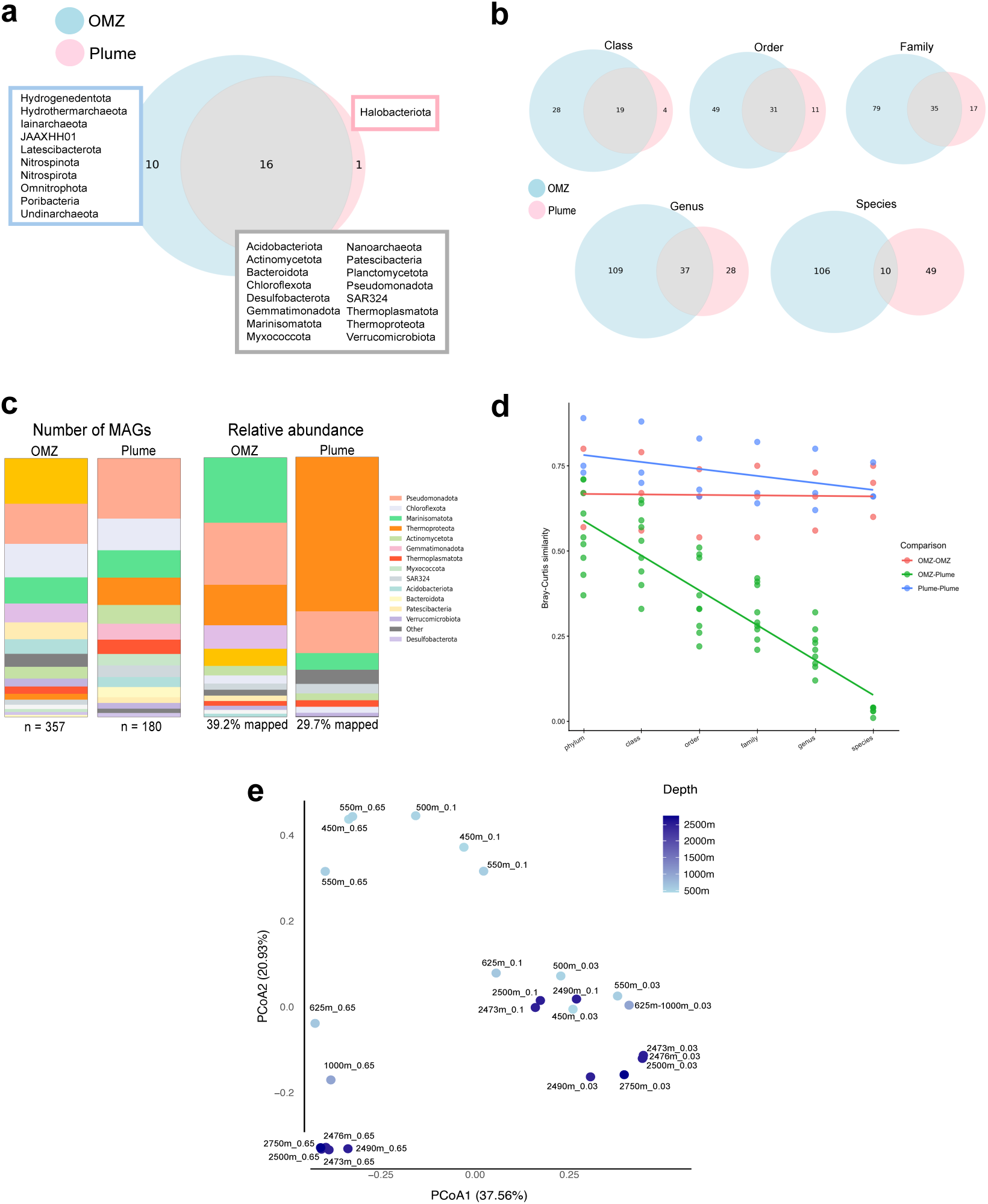
Microbial community differentiation between the plume and OMZ. a) Phyla that are shared or unique to OMZ and plume environments based on recovery of at least one medium-to-high quality MAG from that phylum in the environment b) Number of taxa that are shared or unique to plume and OMZ environments at the class, order, family, genus, and species level based on recovery of at least one medium- to high-quality MAGs from that lineage in the environment. c) Distribution of microbial phyla in the OMZ and plume based on the number of recovered MAGs (left) and their relative abundances (right). The abundance-based bar charts represent only the fraction of assembly reads that mapped to MAGs, as indicated under the respective bar chart. Taxa that comprised <1% of MAG counts or mapped reads were grouped into an “other” category. d) Depth-resolved Bray-Curtis similarity values with varying taxonomic resolution, based on the taxonomy and number of recovered MAGs from each depth. Each point on the graph represents the Bray-Curtis similarity between two individual depths whereas colors indicate if the comparison was within or across environment types e) Principal component analyses of Bray-Curtis similarity values of different samples based on the relative abundance of all MAGs from the dataset, dereplicated at 98% ANI.

Because the taxonomic ranks used for community comparisons can vary (36–38), we assessed taxonomic overlap from the phylum through species levels. Although differences were evident at all ranks, the perceived similarity between OMZ and plume decreased markedly with increasing taxonomic resolution. At the phylum level, 59% of lineages had representatives in both the OMZ and plume (Figure 3a; Supplemental Table 3). The proportion of shared lineages declined progressively from 37% at the class level to 6% at the species level (Figure 3b; Supplemental Table 3). Further, even at the phylum level, differences in the relative contribution of various phyla to total MAGs counts, as well as the relative abundance of respective phyla, were noted between environments (Figure 3c). Across all taxonomic ranks, more lineages were unique to the OMZ than the plume, indicating greater taxonomic richness within the OMZ. Shannon diversity indices, based on MAG taxonomic composition, were slightly elevated in the OMZ as compared to the plume (∼6-15% higher in OMZ) while Simpson diversity indices were more similar (∼0.4-1.8% higher in the OMZ), indicating that the increased richness and diversity in the OMZ is likely driven, in large part, by rare MAGs (39) (Supplemental Table 6).

To assess whether diversification between environments exceeded the variability observed between samples within the same environments, we calculated depth-resolved Bray-Curtis similarity values based on the number and taxonomic composition of MAGs (Supplemental Table 3) and the relative abundances of MAGs across samples (Supplemental Table 7), the latter of which we integrated with principal component analysis (PCoA). Bray-Curtis analyses based on the taxonomy and number of recovered MAGs from each depth demonstrated that, although within-environment similarity remained constant or decreased slightly with increased taxonomic resolution, between-environment similarity decreased more substantially (Figure 3d, Supplemental Table 8). Consistent with these patterns, relative abundance-based PCoA (Figure 3e, Supplemental Table 8) supported both environment type and filter size as major sources of variation, together indicating that strong environmental selective pressure is evident between OMZ and hydrothermal plume microbiomes.

### Taxonomy of individual lineages is an inconsistent indicator of environmental selection

Although the OMZ and plume were consistently distinguishable across all taxonomic ranks at the community level, individual lineages did not always demonstrate the same trends. Some lineages, such as Nitrospinota and Halobacteriota, were environment-specific at the phylum level (Figure 3a), while others diverged only at finer taxonomic levels. Phylum Chloroflexota, for example, was present in both environments but was represented by 17 different families in the OMZ versus only four in the plume (Figure 4a-b; Supplemental Table 3) indicating higher within-phylum taxonomic diversity in the OMZ. Conversely, Marinisomatota showed similar taxonomic diversity but distinct relative abundance patterns (Figure 4a-b; Supplemental Table 3,7). In other cases, taxonomy and abundance alone did not capture environmental differentiation. Phylum SAR324 exhibited identical family level-taxonomy and similar total relative abundance across environments (Figure 4a-b, Supplemental Table 3,7), yet OMZ-associated MAGs encoded additional carbon, nitrogen, and sulfur metabolism genes absent from plume counterparts demonstrating metabolic differentiation (Figure 4c; Supplemental Table 9). This finding aligns with previous reports of depth-dependent metabolic shifts in SAR324 (25). Filtration patterns further suggested environmental differences in size or lifestyle as OMZ SAR324 were recovered on both the 0.65µm and 0.1µm filter fraction but plume SAR324 on the 0.65µm fraction (Figure 4b; Supplemental Table 3).

**Figure 4.**
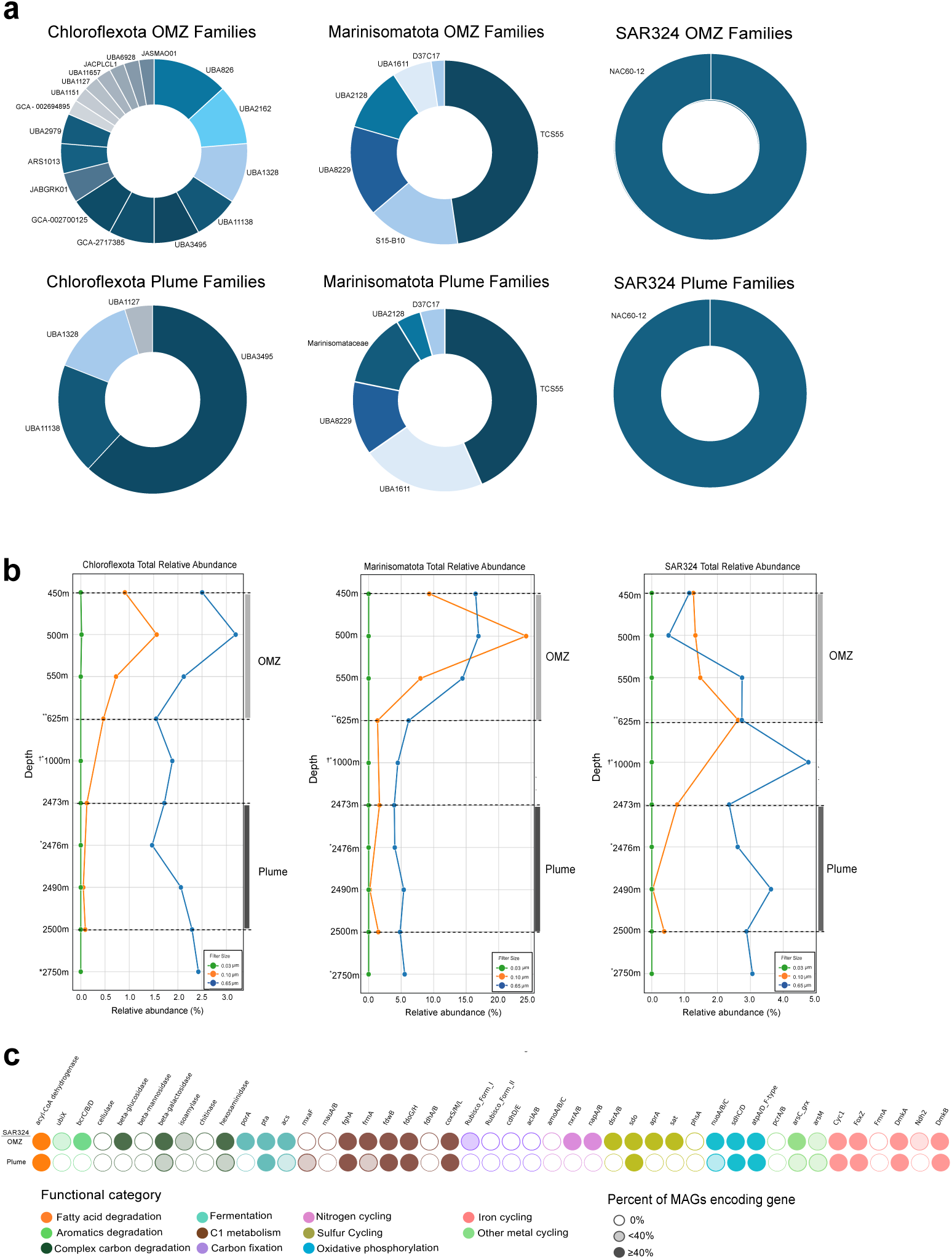
Taxonomy, abundance, and metabolism of select microbial phyla shared between plume and OMZ environments. a-b) Family level taxonomic break down and relative abundance patterns for three different phyla found in both the plume and OMZ. Family-level taxonomic break down was determined using the taxonomy of non-dereplicated MAGs. In the relative abundance plots, * indicates a 0.1µm assembly was not available for that depth, ** indicates a 0.03 µm assembly was not available for that depth, and † indicates that the 0.03µm sample filter was a combination of two depths (625m and 1000m). c) Prescence/absence of key metabolic genes present in the SAR324 MAGs recovered from the OMZ and plume. All SAR324 MAGs recovered in this study are incorporated in the matrix. Genes are color coded by functional category. Comprehensive metabolic profiles are available (Supplemental Table 9).

Together, these contrasting patterns demonstrate that, depending on the taxonomic resolution considered, direct lineage-to-lineage environmental comparisons could either miss the broader, phylum-level restructuring between environments, or overinterpret within-phylum variability as environmental selection when it may reflect intrinsic taxonomic heterogeneity. Accordingly, we additionally integrated relative abundance, metabolic potential, co-occurrence patterns, and insights from our filtration-based size fractionation to identify additional generalizable links between community structure and environmental context, summarized below.

### The OMZ is enriched in smaller microorganisms with limited metabolic capabilities

A clear distinction between the OMZ and plume was the proportion of MAGs recovered from each filter size. In the plume, only ∼15% of MAGs were recovered from the 0.1µm and 0.03µm fractions, whereas >50% of OMZ MAGs originated from these smaller fractions (Figure 5a; Supplemental Table 2), suggesting that the OMZ supports smaller or free-living microorganisms. Supporting this interpretation, MAGs from super-phylum DPANN (Nanoarchaeota, Undinarchaeota, Iainarchaeota) and CPR (Patescibacteria), known for their small cell size, reduced genomes, and symbiotic lifestyles (40–42), were enriched in the OMZ relative to the plume (Figure 5b; Supplemental Figure 4ab; Supplemental Tables 3,7). While DPANN and CPR lineages in OMZs have been previously described (40,43), our study documents a dramatic contrast in both MAG recovery and relative abundance of the groups between the two environments. From OMZ depths, we recovered 67 DPANN and 23 CPR MAGs, whereas only 1 DPANN and 4 CPR MAGs were obtained from the plume. DPANN MAGs were enriched ∼10x in the OMZ (∼2.6% total relative abundance) relative to plume (∼0.24% total relative abundance) environments and CPR were ∼2.5x enriched in the OMZ (∼0.8% total relative abundance) compared to the plume (∼0.3% total relative abundance).

**Figure 5.**
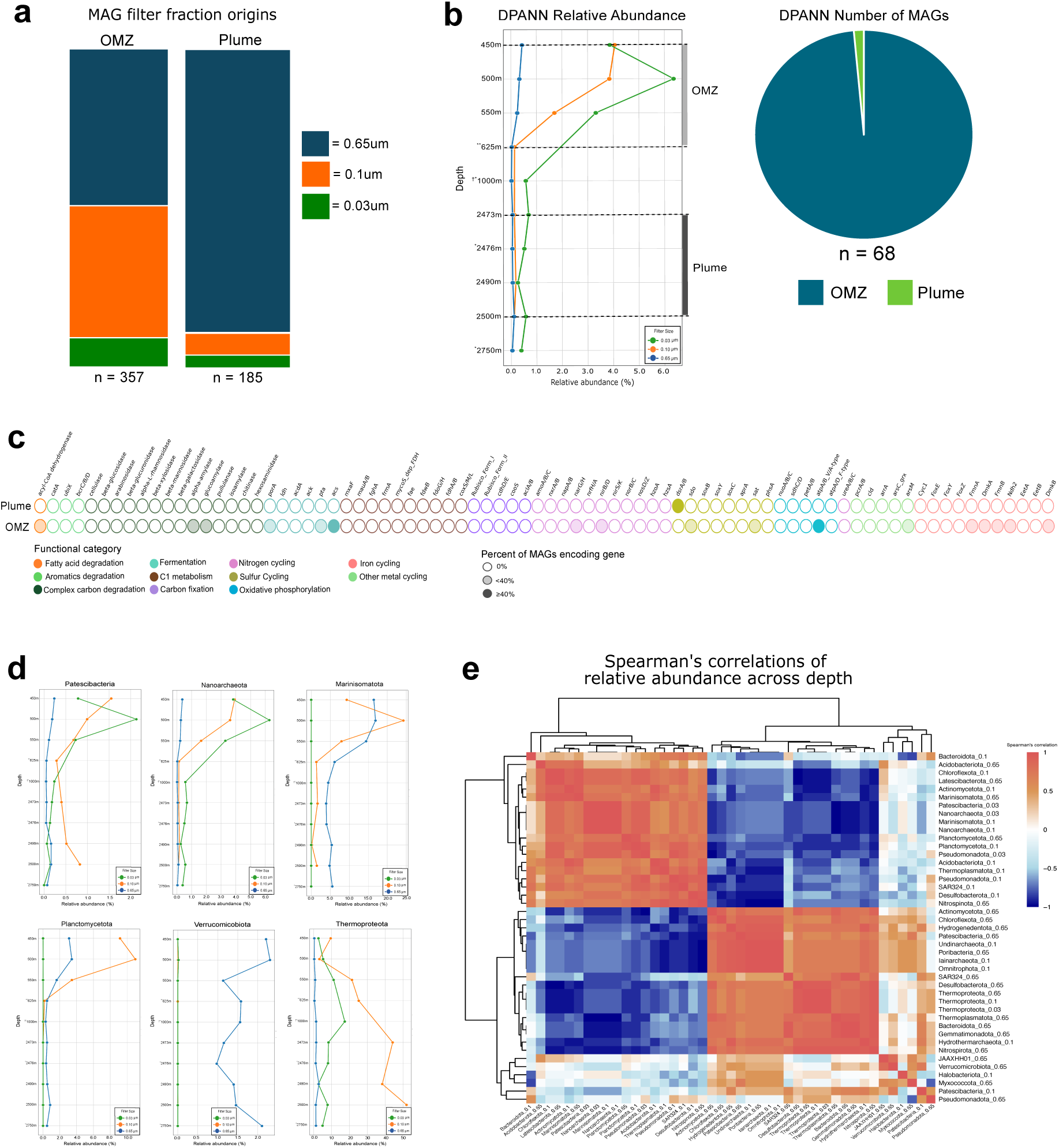
Abundance, metabolism, and co-occurrence patterns of ultra-small bacteria and archaea in the OMZ. a) Break down of the filter fraction from which MAGs were recovered from OMZ and plume environments. The number of MAGs in each environment is indicated next to the respective bar chart. b) Relative abundances of DPANN archaea by depth and filter size. Also included is the breakdown of the environmental origins of DPANN MAGs. In the relative abundance plots, * indicates a 0.1µm assembly was not available for that depth, ** indicates a 0.03µm assembly was not available for that depth, and † indicates the 0.03µm sample filter was a combination of two depths (625m and 1000m). c) Presence or absence of key metabolic genes in DPANN MAGs recovered from the plume and OMZ. d) Examples of microbial phyla whose relative abundance patterns mimic those of CPR and DPANN. Abundance patterns of Acidobacteriota and Thermoproteota, which differ markedly from DPANN and CPR, and included as a reference. e) Heatmap of log-transformed Spearman’s correlation coefficients between all phyla, further separated based on filter-fraction origin.

Consistent with known characteristics (41,44,45), DPANN and CPR MAGs in this study exhibited limited metabolic potential (Figure 5c; Supplemental Figure 4c; Supplemental Table 9). While genes for most components of oxidative phosphorylation were absent from CPR and DPANN genomes, many retained a type A/V ATP synthase, supporting previously documented fermentative capabilities of CPR and DPANN (44).

Energy limited, low oxygen conditions in the OMZ may create favorable conditions for microbes with smaller genome/cell size like DPANN and CPR by reducing resource demands for genome maintenance (46), supporting fermentive lifestyles, and/or by promoting dependence on microbial interactions. In anoxic systems, interactions involving metabolite exchange can be especially important for the process of complex carbon degradation as the complete breakdown of complex organic matter by a microorganism is energetically demanding and may require a partner organism to remove metabolic byproducts, keeping product concentrations low and driving reactions forward (47,48). Given their limited metabolic capacity, fragmented carbon pathways, and well-documented symbiotic tendencies (42,49), the enrichment of DPANN and CPR in the EPR OMZ may be explained by participation in syntrophic carbon degradation (50). Supporting this interpretation, DPANN and CPR are enriched not only in the OMZ relative to the plume, but also display higher relative abundances in shallower OMZ waters that declined with depth (Figure 5b; Supplemental Figure 4a, Supplemental Table 7), consistent with reliance on photic-zone derived carbon that would become progressively re-mineralized with depth (52). Many CPR and DPANN MAGs also encoded iron-reduction-associated genes (*dmkA*, *dmkB*, *fmnA*, *ndh2*) (Figure 5c; Supplemental Figure 4c; Supplemental Table 9) implicated in flavoprotein-dependent electron transport (53). Given prior observations that DPANN and CPR can be physically associated with host cells (45,54), reducing equivalents may transfer directly between partners, enabling membrane-associated iron reduction and redox balancing within symbiotic partnerships.

Relative abundance patterns further implicated specific microbial partnerships in the OMZ (Figure 5d; Supplemental Figure 5). Spearman’s correlations of log-transformed relative abundances revealed strong positive associations (r > 0.80) between Nanoarchaeota, the dominant DPANN phylum, and taxa such as Marinisomatota and Planctomycetota. CPR similarly correlated with phyla including Nitrospinota, Chloroflexota, and Desulfobacterota (Figure 5e; Supplemental Table 10). Notably, metabolite sharing has previously been suggested to be a strategy of abundant OMZ phylum Marinisomatota (55), further pointing to this lineage as a potential symbiotic partner of DPANN and/or CPR. Together, these results suggest that OMZ conditions select for smaller cell size, metabolic streamlining, and potentially cooperative metabolic exchange.

### Availability of redox substrates drives several community-wide metabolic shifts

Microbial metabolism and abundance are often strongly influenced by environmental geochemistry (56–58). To assess how environmental pressures shape microbial communities, we quantified the relative abundance of genes involved in nitrogen, carbon, sulfur, metal, and mineral metabolism in the OMZ and the plume (Figure 6a-d; Supplemental Figure 6) using both MAGs and un-binned assemblies to account for methodological biases

**Figure 6.**
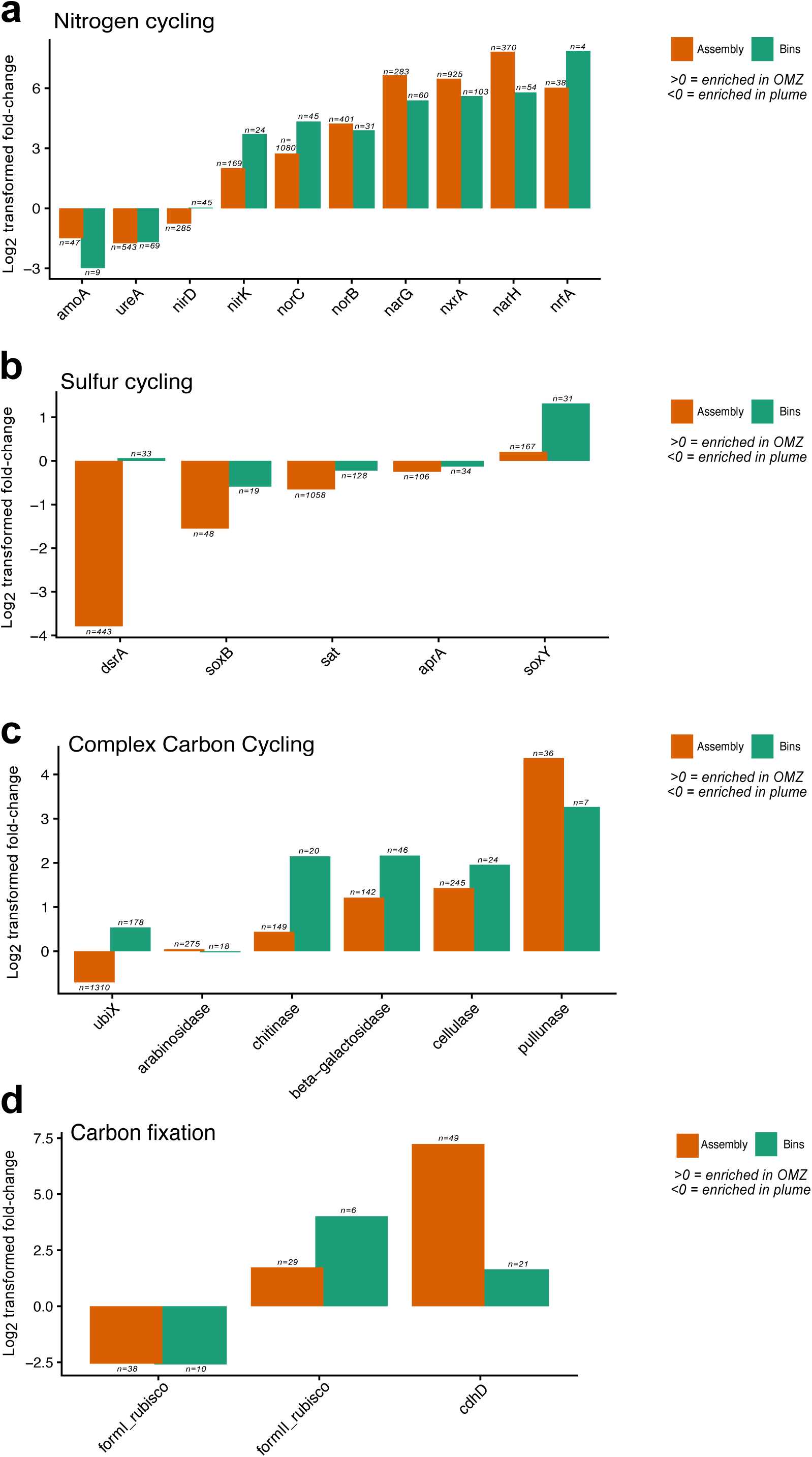
Relative enrichment of various metabolic genes in the OMZ and plume. a-d) The log_2_ transformed value of the abundance fold change between the OMZ and plume environments for various gene categories are shown. Values were calculated based on relative abundance values of MAGs encoding the genes (MAGs) or RPKM values of gene encoding scaffolds from un-binned assemblies (assemblies). Values greater than zero indicate the gene was enriched in the OMZ as compared to the plume whereas values less than zero indicate the gene was enriched in the plume as compared to the OMZ. The number of genes or MAGs used for each calculation is indicated above the respective bars in the figure. Acronyms: *amoA* = methane/ammonia monooxygenase subunit A; *ureA* = urease subunit gamma; *nirD* = nitrite reductase small subunit; *nrfA*= nitrite reductase cytochrome c-552; *nirK*= nitrogenase molybdenum-iron protein beta chain; *norC*= nitric oxide reductase subunit C; *norB* = nitric oxide reductase subunit B; *nxrA* = nitrate reductase / nitrite oxidoreductase, alpha subunit; *narG*=nitrate reductase / nitrite oxidoreductase, alpha subunit; *narH*=nitrate reductase / nitrite oxidoreductase, beta subunit; *ubiX* = flavin prenyltransferase, pullunase; *cdhD* = CODH/ACS complex subunit delta K00194; *dsrA* = dissimilatory sulfite reductase alpha subunit; *aprA* = adenylylsulfate reductase, subunit A; *soxB* = S-sulfosulfanyl-L-cysteine sulfohydrolase; *soxY* =sulfur-oxidizing protein; *sat* = sulfate adenylyltransferase.

MAG-based analyses revealed enrichment of genes for nitrogen reduction processes (*nar, nor, nrf, nir, nxr)* (59) in the OMZ (Figure 6a; Supplemental Table 11). This pattern is consistent with nitrate serving as a favorable electron acceptor under oxygen-limited conditions and previous reports of substantial microbial nitrogen cycling in OMZs (14,60). Additionally, MAGs from phyla Nitrospinota and Nitrospirota, known for their nitrogen cycling capabilities (61), were recovered exclusively from the OMZ (Figure 3a; Supplemental Table 3). Similarly, several genes for the reduction of iron, an additional strong electron acceptor, were enriched in the OMZ as compared to the plume (Supplemental Figure 6). Conversely, genes for urea utilization and ammonium oxidation were enriched in the plume, postulated to be a result of increased oxygen availability, or symbiotic partnerships with eukaryotes that frequently reside at vents and generate urea as a waste product (62) (Figure 6a; Supplemental Table 11). Enrichment of phylum Thermoproteota, a known ammonium oxidizing archaeal phyla (62,63), was also noted in the plume (Figure 3c; Supplemental Figure 5). Genes for the metabolism of manganese and chlorite were also slightly enriched in the plume likely due to inputs of these minerals from the hydrothermal vent source (Supplemental Figure 6, Supplemental Table 11) (64,65).

Microbial dissimilatory sulfur oxidation and/or reduction has also been implicated to occur in both hydrothermal plumes and OMZs (66,67). Indeed, genes for dissimilatory sulfur reduction/oxidation (*dsrA*, *aprA*, *sat*, *soxY, soxB*) (68) were present in both environments and several showed a slight enrichment in the plume consistent with sulfide inputs from venting sources (Figure 6b; Supplemental Table 11). However, no MAG encoded a complete canonical sulfate reduction pathway (*sat*-*apr*-*dsr*) in either environment (Supplemental Table 9). The presence of oxidative *dsr* genes *(dsrEFH),* and/or the absence of the reductive marker gene (*dsrD),* suggest that *dsr-*encoding MAG*s* in our dataset were of the oxidative type (69,70). While partial dissimilatory sulfate reduction via *sat* and *apr* may be possible, full sulfate reduction does not appear to be an energy generating strategy in either environment, contrasting with previous reports of substantial sulfate reduction potential in OMZs and hydrothermal systems (66,67,71).

Genes involved in complex carbon degradation were largely enriched in OMZ MAGs relative to plume MAGs, potentially reflecting high organic matter inputs from surface waters (Figure 6c; Supplemental Table 11). Both OMZ and plume MAGs encoded carbon fixation genes however, the pathways differed between environments (Figure 6d, Supplemental Table 11). OMZ MAGs were enriched in the Wood-Ljungdahl pathway associated gene *cdhD* and the Calvin-Benson-Bassham (CBB) cycle associated gene, form II Rubisco, whereas plume MAGs lacked Wood-Ljungdahl pathway gene *cdhD* and were enriched in genes associated with form I Rubisco-mediated carbon fixation (Figure 6d; Supplemental Table 9,11). These differences likely reflect pathway optimization based on environmental redox conditions (72–74) but demonstrate that carbon fixation does occur well into the dark ocean, a phenomenon that has been increasingly documented to impact oceanic carbon burial and storage (75,76).

For most genes, the directionality of enrichment patterns between the OMZ and plume was consistent across both MAG-based and assembly-based analyses (Figure 6a-d; Supplemental Table 11-12) however, several genes exhibited notable method-dependent discrepancies. For example, the dissimilatory sulfite reduction/ sulfur oxidation gene *dsrA* appeared invariant between environments in MAG-based analyses, yet was ∼5x fold enriched in plume samples when assessed using assemblies. Similarly, carbohydrate-active gene *ubiX* was enriched in the OMZ in MAG-based analyses, but showed plume enrichment in assembly-based results. Such discrepancies may reflect incomplete binning in MAGs or gene redundancy in assemblies, but the contrasting enrichment directions warranted further investigation.

### Viral-encoded metabolic genes can contribute significantly to community-wide patterns

We sought to understand the method-based discrepancies seen in some enrichment patterns. Examination of the scaffolds underlying genes such as *dsrA* demonstrated that many high coverage scaffolds encoding the gene of interest originated from plume-derived 0.1µm and 0.03µm filter fractions (Supplemental Table 11), despite low MAG recovery from these size fractions in the plume. Given the documented viral enrichment in the 0.03µm (Supplemental Figure 3d) fractions and prior evidence that deep-sea viruses can carry metabolic genes as auxiliary viral genes (AVGs) (77–79), we investigated whether these scaffolds had viral origin.

We screened all scaffolds encoding the 33 examined metabolic genes using viral identification software and found that 6 genes were encoded on a viral-predicted scaffold in at least one instance (Figure 7a; Supplemental Table 13). Notably, two genes displaying method-dependent discrepancies including, *dsrA* and *ubiX*, were amongst the genes encoded on viral-associated scaffolds. *DsrA*-encoding scaffolds were particularly enriched in viral predictions, with 30 of the 474 *dsrA*-encoding scaffolds (∼6%) identified as viral (Supplemental Table 13). Sulfur-cycling gene encoding phages have been noted in plume microorganisms previously (78,79) but our results suggest that interestingly, in the case of *dsrA*, this may be a phenomenon specific to hydrothermal vent environments as *dsrA* encoding viral scaffolds were not detected in the OMZ. In contrast, *soxY* was associated with viral scaffolds only in the OMZ which may explain why *soxY* showed enrichment in the OMZ, a pattern opposite from other sulfur genes.

**Figure 7.**
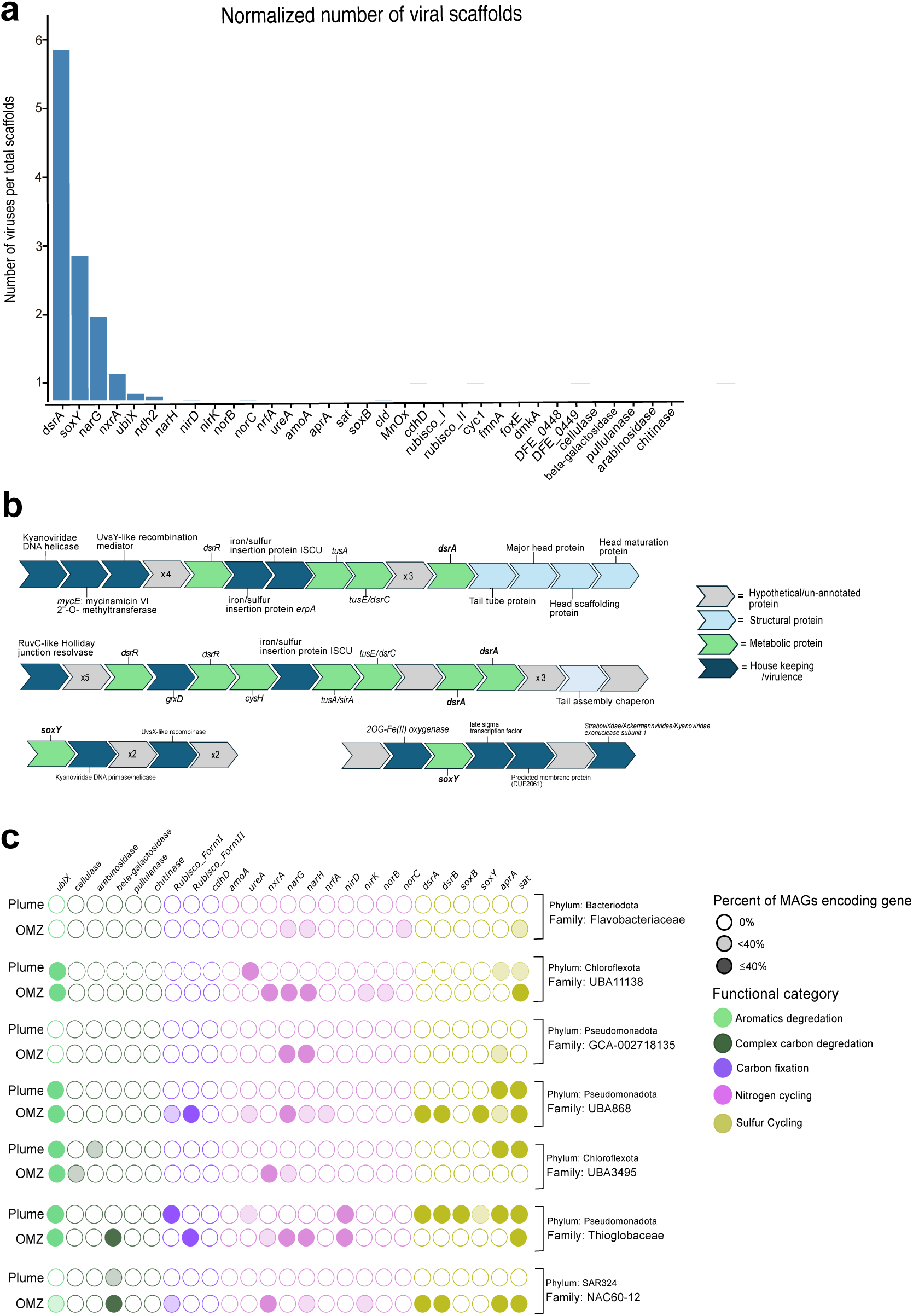
Viral metabolic genes and lineage specific metabolic shifts at EPR. a) Number of gene-encoding scaffolds predicted to be of viral origin, normalized to the total number of scaffolds encoding the gene. b) Genomic architecture of select plume derived *dsrA* and *soxY* encoding scaffolds predicted to be viral. c) Lineage specific metabolic shifts in nitrogen, carbon, and sulfur metabolism.

We further validated viral involvement by examining the genomic architecture of representative *dsrA and soxY*-encoding scaffolds with viral predictions (Figure 7b). These regions exhibited clear viral genomic organization, including genes encoding capsid, tail, and replication-associated proteins, as well as recombinases (77), supporting a viral rather than cellular origin. In some cases, we could verify viral genes flanking the metabolic gene on only one side, raising the possibility that the gene belongs to the bacterial host of a prophage and was incorrectly assembled with the virus. However, the observed genomic architecture, the filter fraction in which the genes were recovered, and the discrepancy in abundance patterns provides strong preliminary evidence of environment specific viral involvement in metabolic processes.

### Lineage specific metabolic shifts may differ from community wide patterns

In addition to investigating metabolic genes within the community, we investigated gene patterns within specific lineages to determine whether the observed metabolic shifts occurred uniformly across the community or if they were lineage specific (Figure 7c). We examined seven family-level lineages for which at least three MAGs were recovered from both the plume and the OMZ. For nitrogen and carbon cycling, most lineages mirrored the trends observed at the community level. For example, in lineage UBA1138, MAGs originating from the OMZ encoded one or more of *nxr* (nitrite oxioreducatase) *narG/H* (nitrate reductase) or *norB/C* (nitric oxide reductase) while these genes were completely absent from plume counter parts. In turn, plume counter parts encoded *ureA* which was absent from OMZ MAGs, exactly following the community wide patterns. Similarly, in the majority of cases, OMZ lineage counter parts showed higher frequency of MAGs encoding complex carbon degradation genes.

Some gene patterns, however, showed less consistency across lineages. Some lineages, such as Pseudomoadota family *Thioglobaceae*, followed community-wide patterns, encoding sulfur cycling genes in the plume that were absent in the OMZ. Other lineages, such as SAR324, displayed contrasting patterns (Figure 7c). Likewise, MAGs from lineage UBA868 encoded both forms of Rubisco in the OMZ, but none in the plume, also digressing from community patterns. These findings emphasize that while certain environments may broadly favor specific metabolic processes, individual lineages do not always conform to these trends. Factors such as metabolic flexibility, competitive pressures from other community members, or susceptibility to viral influence likely shape how a given lineage allocates its metabolic repertoire (80).

## DISCUSSION

The importance and complexity of microbial communities on a changing planet is increasingly well recognized (4) however, substantial knowledge gaps remain regarding how environmental microbial communities are structured, how they evolve, and how their metabolic interactions respond to environmental change. These gaps limit our ability to predict ecosystem responses to climate change and to evaluate the risks and benefits of proposed environmental intervention strategies (81–83). In this study, we generate over 700 novel MAGs from the deep sea, not only expanding the catalog of microbial diversity in understudied extreme environments but also helping to disentangle the interacting ecological and thermodynamic factors that shape and sustain microbial communities in the ocean.

Through integrated metagenomic analyses, we demonstrate that although microbial communities in OMZs and hydrothermal plumes can appear broadly similar at coarse taxonomic resolution, analyses at higher taxonomic levels reveal clear environmental structuring. These findings highlight that the concept of taxonomic sufficiency – often used in macroecological studies to determine what is the lowest resolution of taxonomy needed to meet the goals of a particular study (84) – is also relevant to microbiology studies, playing a major role in analytical methods and interpretation of findings. By combining taxonomy with investigation of other parameters, such as relative abundance, metabolic potential, and size fractionation, we reveal additional pronounced and ecologically meaningful differences. Our findings indicate that thermodynamic constraints imposed by redox conditions, symbiotic relationships among microbes, as well as viral predation are potential major evolutionary drivers structuring these communities (Figure 8a-f). Shifts in these parameters may therefore be likely to result in disproportionately large changes in microbial community structure and function.

**Figure 8.**
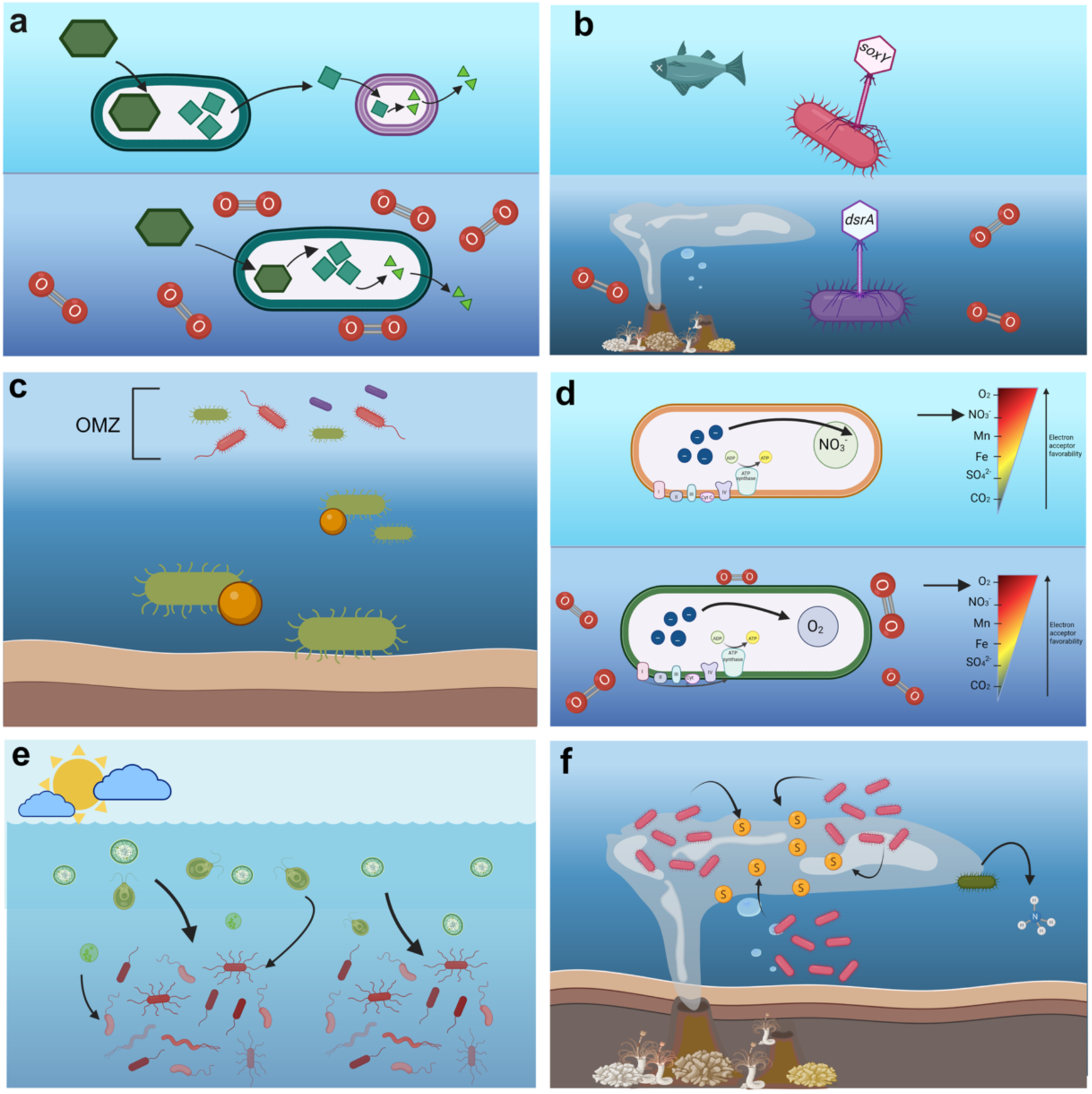
Schematic of proposed selective pressures structuring plume and OMZ microbial communities. a) Thermodynamic constraints imposed by low oxygen concentrations in the OMZs favor fragmented carbon degradation pathways and the presence of small auxotrophic microbes that participate in the exchanges. b) Viral-encoded metabolic genes such as *dsrA* and *soxY* can account for a large proportion of the gene abundance in the plume and OMZ that is not reflected by examining microbial genomes alone. c) General trends toward increased taxonomic richness and reduced genome size, possibly due to low-energy, oxygen limited conditions, are observed in the OMZ. d) Depletion of oxygen in the OMZ causes microorganisms to develop metabolisms that use alternate electron acceptors such as nitrate. e) High influxes of organic matter into the photic zone lead to an increase in genes for carbon degradation in microbial MAGSs. f) Individual lineages may change metabolic strategies based not just on the most favorable redox substrate but also on the relative competition for the specific metabolite. Figure generated with Biorender.com

Our findings also have implications for broader microbiology applications and problems beyond marine ecology. For example, members of the DPANN archaea have been characterized as “parasitic” or “vampire”-like, reflecting their unidirectional metabolic dependence – and often detrimental effects – on other community members (49,85). Our analyses suggest these organisms may also participate in essential commensal relationships, exerting influence not primarily through direct metabolic exchange, but through thermodynamic regulation, driving forward complex carbon degradation reactions by consuming inhibitory by-products in oxygen limited environments. Given that most environmental microbes remain difficult to culture (86,87) these results underscore the need to rethink cultivation strategies not only in terms of substrate provision but also with respect to environmental constraints and inhibitory conditions that may be mitigated only through complex community interactions.

Further, while this study emphasizes contrasts between OMZ and hydrothermal plume communities, it is equally important to note the presence of shared microbial lineages across both environments, in some cases extending to the species level. These shared taxa are likely characterized by substantial metabolic and physiological flexibility, including tolerance to variable oxygen concentrations, temperatures, pressures and depths. Such traits point to these lineages as promising candidates for future bioengineering or synthetic ecology applications. Although metabolic flexibility in microorganisms is often viewed as advantageous for environmental engineering applications (88,89), it may also present an underappreciated limitation. In dynamic systems, this flexibility could lead to incomplete or transient remediation, as microbial communities shift their functional priorities once selective pressure decreases. Thus, the same resilience that enables microorganisms to survive environmental perturbations may also undermine the stability and predictability of engineered bioremediation outcomes.

Further while this work provides new insights into microbial diversity and selective pressures associated with hydrothermal plumes and OMZs at the East Pacific Rise, the extent to which these patterns generalize across the global ocean remains uncertain. Although OMZs and hydrothermal systems worldwide share defining characteristics, they vary considerably in spatial extent, geographic location, and geochemical context (90,91). Accordingly, our findings should be viewed as a conceptual framework for interpreting microbial community differentiation rather than a universal set of rules. Extending similar integrative analyses across diverse oceanographic settings will be critical for assessing the broader applicability of these patterns.

Finally, our study provides an overview of two environments that differ in many parameters including depth, temperature, and geochemistry. While we are able to postulate which parameters drive which changes in the microbial communities based on the observed change and known properties of microorganisms, disentangling exactly which parameter is driving community divergence with confidence remains difficult. Future studies would greatly benefit from integrating additional data points as references.

## METHODS

### Sample collection

Samples were collected aboard the *R/V Atlantis* in December of 2018 on cruise AT42-06. Briefly, a conductivity, temperature, and depth (CTD) sensor mounted on a rosette containing 24 12-liter Niskin bottles was deployed on a vertical cast. The Niskin bottles were closed at depths in the water column corresponding to signals which dictated hydrothermal plumes, OMZs, or background water. The presence of a hydrothermal plume was indicated by an anomaly in temperature and/or turbidity at depths and locations of known venting sites while an OMZ was dictated by oxygen concentrations below 20 micromolar (Supplemental Figure 1). After recovery, microbial communities were collected through sequential serial filtration of seawater first through a 0.65µm filter, followed by a 0.1µm and 0.3µm filter, using Harrington Industrial LLC brand filters (ZTECG 0.65-5P3S, 5” WATERTEC 0.1-5P3S, ZTECE 0.03 5P3S) and a peristaltic pump. Immediately after sampling, filters were soaked with RNA later (14.3 mM ethylenediaminetetraacetic acid, 17.9 mM sodium citrate trisodium, 3.78 M ammonium sulfate) for preservation and stored at -80 °C until further processing. Additional information on sampling sites, including geographic coordinates are included (Supplemental Table 1). For one of the 0.03µm samples, two depths (625m and 1000m) were combined on the same filter. This is indicated in figures and metadata as needed.

### DNA extraction and library preparation

Genomic DNA was extracted from filters using a phenol/chloroform extraction followed by ethanol purification. Filters were first removed from their plastic casing using an ethanol sterilized rotating saw. A section of filter was briefly thawed and removed using sterilized scissors and tweezers. The outer layers of the filter were removed, and the filter was cut into small pieces (approximately 1sq cm in size). Filter pieces were placed into a 50ml sterile conical tube. If not immediately proceeding with the next steps of the protocol, the filter and tube were stored at -80°C. 7ml of pre-made DNA Extraction Buffer (0.1M Tris, 0.1M EDTA, .05M NaH2PO4, 0.3M NaCl), 140 µl of Lysozyme (50mg/ µl), and 350 µl of 1M DTT (dithiothreitol) were added to the tube containing the filter. The tube was vortexed and inverted to distribute buffer and reagents, placed on a tube rotator (25rpm), and incubated at 37 °C for 30 minutes. After incubation, 180µl of proteinase K (20mg/µl) and 350µl SDS (20%) were added to the tube containing the filter. The tube was vortexed and inverted again to distribute the reagents and placed back on the rotator to incubate at 65 ° C for 45-90 minutes. Reagent only negative controls were included intermittently during the DNA extraction process.

After incubation, liquid from the 50ml conical tube was decanted into a sterile 15ml polypropylene conical tube containing approximately 1ml of autoclave sterilized silica/zirconia beads (sizes 0.1mm, 0.5mm, 1mm). The tubes were bead beat (6m/s, 60s, 2 cycles, model MP-Bio FastPrep25-5G), then incubated on ice for 10-15 minutes. After incubation on ice, tubes were centrifuged (2,500xg for 10 minutes) to pellet the beads and cell debris. The supernatant was decanted into a new 15ml falcon tube, and an equal volume of 25:24:1 phenol chloroform isomyl alcohol (pH 8) was added. Tubes were inverted repeatedly to mix, and centrifuged (2500xg, 10 minutes) to separate the aqueous and organic layers. The aqueous layer was removed from the 15ml tube with a pipette and placed into a new 15ml sterile conical tube. Equal volumes of 24:1 chloroform:isoyml alcohol were added, and the tube was again inverted to mix and subsequently centrifuged at (10 min, ∼2500xg). After centrifugation, the aqueous phase was removed with a pipette and placed into a sterilized, 50ml round bottom centrifuge tube. Two volumes of chilled, 100% molecular grade ethanol, and a 1/50 dilution of glycogen (20mg/µl) were added to the tube. The tubes were gently rolled to mix then incubated at -20°C for 0.5-2 hours, or overnight.

After incubation, the samples were centrifuged (15,000 x g, 30 minutes) to pellet the DNA. After centrifugation, the supernatant was removed, replaced with 10ml, chilled, 70% molecular grade ethanol, and centrifuged (15,000 x g, 30 minutes) The supernatant was removed, and the tubes were inverted in a biological safety cabinet to air dry. The pellets were resuspended by adding sterile, nuclease free water and incubated at 50°C in a water bath for 5-10 minutes until the pellet was redissolved. For some samples which failed initial sequencing efforts, DNA was purified using a 3k omega Nanosep centrifugal filtration device to remove excess salts (VWR, 29300) Briefly, the column was first sanitized by washing with 500µl of 70% ethanol and centrifuging at 13,000xg. The column was rinsed twice by adding 500µl sterile water and centrifuging. The column was used within 20 minutes to avoid drying. The DNA sample was eluted in sterile nuclease free water, placed in the centrifugal column, and centrifuged for 5-10 minutes at 13,000 x g. The process was repeated until all the sample was passed through, and the resulting eluted DNA was pipetted off from the filter. The DNA concentration was determined using a Qubit^TM^ 1X HS Assay Kit (Q33230). Library prep on purified DNA was carried out by the U.S Department of Energy Joint Genome Institute (92) using either Illumina ss/dsDNA-Seq kit or Illumina Low Input Fragment for samples with very small DNA concentrations.

### DNA sequencing and assembly

DNA was sequenced using Illumina short read technology at 2×151bp at the U.S Department of Energy Joint Genome Institute (JGI) (92). Assembly of reads was also accomplished at JGI with the following methods: Reads were trimmed and adapter and contaminant sequences removed using BBDuk (93). For assembly, reads were further corrected using BBtools v38.90 (93) with the following command: ‘-Xmx100g metadatafile=counts.metadata.json mincount=2 highcountfraction=0.6 in=bbcms.input.fastq.gz out1=input.corr.left.fastq.gz out2=input.corr.right.fastq.gz.’ Corrected reads were assembled using metaSPADES v3.15.2 (94) with the following command: ‘-m 2000 --tmp-dir cromwell_root -o spades3 --only-assembler -k 33,55,77,99,127 --meta -t 16 -1 input.corr.left.fastq.gz -2 input.corr.right.fastq.gz’

### Metagenomic binning and viral identification

Binning of microbial genomes was accomplished using metaWRAP v1.3.2 (95) with the ‘Initial Binning’ and ‘Bin Refinement’ modules. The initial binning module was run with MaxBin (96) and MetaBAT2 (97) modules and the ‘--universal’ flag to improve archaeal binning. The bin refinement module was run with default parameters and the ‘-c 50 -x 10’ flags to select for only medium-to-high quality bins (32) (≥50% competition and ≤10% contamination). Each assembly was binned individually and utilized non-corrected, filtered reads from all depths and filter sizes to improve differential coverage profiles. Viral scaffolds from each assembly were identified by geNomad v1.7.4 (98) utilizing the first module of the MVP v1.1.4 viral wrapper with default parameters (99). To visualize viral genomic structure, CheckAMG (https://github.com/AnantharamanLab/CheckAMG) was run on viral scaffolds and the resulting annotations were used to manually reconstruct the gene regions of interest.

### Taxonomic classification and metabolic analysis

Taxonomic classification of MAGs was performed with GTDBtk v2.4.0 (100) using the r220 database with the “classify_wf” option. Metabolic analyses and functional annotation of MAGs for constructing presence/absence figures was conducted using METABOLIC-G v4.0 (101) on DNA sequences using default parameters. The presence/absence of various genes was determined based on the results of the METABOLIC HMM search function (result worksheet 1).

### MAG quality assessment and dereplication

The completion and contamination of MAGs was evaluated using CheckM 1.0.12 (102) as part of the MetaWRAP ‘Bin Refinment’ pipeline. The completeness and contamination of resulting MAGs was verified using an independent run of CheckM 1.2.0. Completeness and contamination values for CPR (Patescibacteria) were evaluated using a custom set of gene markers, tailored specifically for the reduced genome size of the phylum (103). In some cases, evaluation of Patescibacteria MAG quality using custom markers demonstrated that MAGs exceeded contamination thresholds, however the genomes were retained due to previous calculations from MetaWRAP and the fact that their small size may skew contamination thresholds higher. To calculate estimated genome size, the MAG size in base pairs was divided by the estimated genome completion.

For analyses requiring genome dereplication, genomes were dereplicated using dRep v.3.3.0 (104) with the following flags ‘-nc 0.5 -con 10 -comp 50.’ Genomes were dereplicated at 95% ANI for species level analysis and 98% ANI to generate a non-redundant set of bins for read mapping and relative abundance calculations, described in more detail below.

Because low-biomass environmental samples are particularly prone to misinterpretation due to external microbial contamination during the sampling, DNA extraction, and/or sequencing procedures (105), we included multiple reagent-only negative controls alongside DNA extractions to help detect potential DNA contamination. DNA yields from the negative controls were consistently low, ranging from undetectable (“out of range” on the Qubit fluorometer (Thermofisher)) to a maximum of 0.1 ng/μL. Four negative controls with nonzero concentration readings underwent library preparation and sequencing in parallel with the environmental samples. Of these, two samples did not produce assemblies that passed quality control thresholds. The remaining two yielded assemblies which underwent binning and taxonomic classification.

Binning was performed on the negative control assemblies using both negative control read sets and read sets from all other samples to detect external and cross-contamination. Two bins were produced from the negative control assemblies and assigned taxonomy, as described above. Bins with these taxonomic classifications were not present in the environmental data set and thus required no additional curation. In addition, reads from negative control samples were mapped to all bins obtained from the environmental dataset, with mapping detected for only 1 of 700 bins. This bin was subsequently removed from downstream analyses. Additionally, we searched previously published literature that has sought to identify microbial taxa that are common laboratory contaminants (105,106). MAGs from our dataset affiliated with these groups were evaluated using multiple criteria, including taxonomic assignment, sequence similarity, and abundance patterns. Six MAGs classified within the families Enterobacteriaceae and Xanthomonadales were removed from analyses due to (i) taxonomic affiliation with known contaminant groups, (ii) high sequence similarity to non-marine or human-associated microorganisms based on BLAST searches (Supplemental Table 14), and (iii) inconsistent abundance profiles across samples.

For downstream analyses, contigs from contaminant bins were removed from the respective assemblies using assemblies using SeqKit2 (107). To remove contamination from reads, all reads were mapped to the reference contaminant bins and contaminant reads were removed using bbmap (93) with the “minid=0.98” flag.

### Taxon-level relative abundance estimations

To generate a non-redundant MAG set for relative abundance estimations, medium- to high-quality MAGs from all depths and filter sizes were dereplicated at 98% ANI using dRep. Relative abundances of dereplicated MAGs across all 26 metagenomes were calculated using CoverM v0.7.0 (108) (genome mode, default parameters) using filtered, contaminant removed, reads. Taxon-level relative abundances were obtained for each sample by summing the relative abundances of all MAGs assigned to a given taxon.

To estimate a consolidated taxon-level relative abundance within each environment, we first calculated a “per-environment” relative abundance for each MAG by averaging its relative abundance across all samples within that environment. MAG-level abundances were then summed within each taxonomic group, as above, to yield environment-level taxonomic abundances. To ensure comparability between environments, estimations for consolidated environmental abundances were restricted to three depths per environment for which all filter sizes were successfully sequenced (OMZ: 450m, 500m, and 550 m; plume: 2473m, 2490m, and 2500 m). This ensured that each environment-level estimate was derived from the same number of samples with identical filter-size representation.

To estimate the relative contribution of viral sequences to each assembly, filtered, contaminant-removed, reads were mapped to identified viral scaffolds using CoverM (genome mode), and the percentage of mapped reads was reported for each assembly.

### Gene abundance estimations

To estimate gene abundances in each environment using MAGs, we first identified open reading frames using prodigal (109) and identified all MAGs encoding a given gene with hmmsearch (110), applying the trusted cutoff (TC) thresholds established in METABOLIC. HMM profiles were taken from METABOLIC unless a more recent version was available, in which case the newer profile was used. The full set of HMM profiles utilized for these analyses is available at https://github.com/AnantharamanLab/Microbial-Ecology-of-East-Pacific-Rise. We then summed the estimated per-environment relative abundances (described above) of all MAGs encoding the gene. Multiple gene copies within a MAG were not considered in these analyses.

To estimate gene abundances independently of binning, we predicted genes of interest directly from the contaminant-removed assemblies using the same approach. We extracted the contigs encoding these genes and calculated their coverage with CoverM (contig mode, RPKM (reads per kilobase of contig per million mapped reads) method), using contig coverage as a proxy for gene coverage. The total abundance of each gene in an environment was calculated by summing the coverage of all genes originating from assemblies of that environment. In the case that more than one HMM profile was available for a gene, all profiles were used, results consolidated, and redundant scaffolds removed. As above, these analyses were restricted to three depths per environment with complete filter-size representation.

### Phylogenetic analyses

Phylogenetic trees were constructed as previously described (111). Briefly, open reading frames from each MAG were predicted using prodigal, and HMMsearch was used to identify 16 ribosomal protein subunit sequences using either bacterial or archaeal reference proteins. Sequences were aligned using MAFFT (112) and uploaded to Geneious (113) for manual curation where fragmented sequences were removed. The alignments were concatenated, and only concatenated sequences that were > ∼60% the length of the consensus sequence length were retained. The sequences were then re-aligned in MAFFT and the alignment was masked at 95% in Geneious to remove gaps. The phylogenetic tree was constructed using IQ-TREE (114) with the following parameters ‘-T AUTO -ntmax 30 -bnni -bb 1000 -m TESTMERGE’ and visualized using iToL (115). Ribosomal protein alignments and concatenated tree files are included (Supplemental Files 1-4).

### Statistical analyses and data visualization

Statistical analyses and data visualization were conducted using custom scripts in R and/or Python. Bray-Curtis similarity based on MAG taxonomy (Figure 3d) was calculated using a matrix consisting of the number of MAGs obtained from each taxonomic lineage at each depth. Bray-Curtis similarity based on MAG abundance (Figure 3e) was calculated using a matrix of the relative abundances of each MAG across each assembly, using the set of MAGs dereplicated at 98% ANI. Spearman’s correlation coefficients (Figure 5e) were calculated based on a matrix of the total relative abundance of each phylum from each depth with each phylum further delineated by filter fraction. For example, “Actinomycetota_0.65” consisted of the summed relative abundances for all Actinomycetota MAGs originating from the 0.65 filter fraction whereas “Actinomycetota_0.1” consisted of the summed relative abundance of all Actinomycetota from the 0.1 filter fraction. All scripts for statistical analyses and data visualization are available at (https://github.com/AnantharamanLab/Microbial-Ecology-of-East-Pacific-Rise). Included with each script are the input data. Generative AI (OpenAI) was used for assistance with generating and de-bugging scripts. The scripts and their outputs were manually inspected and validated.

## Supporting information

All supplemental tables associated with the manuscript

Alignments and tree files for phylogenetic trees in the manuscript

## Acknowledgements

K.A. acknowledges support by the National Science Foundation under grant numbers DBI2047598 and OCE2049478. The authors also acknowledge support under proposal 505680 (10.46936/10.25585/60001283) by the U.S. Department of Energy Joint Genome Institute (https://ror.org/04xm1d337), a DOE Office of Science User Facility, supported by the Office of Science of the U.S. Department of Energy operated under Contract No. DE-AC02-05CH11231. J.C.K. was supported by a National Science Foundation Graduate Research Fellowship. We would also like to acknowledge Woods Hole Oceanographic Institution and the captain and crew of the *R/V Atlantis* and HOV *Alvin* for their assistance in sampling.

## DATA AVAILABILITY

Raw data (reads, assemblies, metagenome assembled genomes) are made available to reviewers via Dryad. Data has been submitted to the European Nucleotide archive under PRJEB128667 and will be submitted for public release following peer review and publication.

## SUPPLEMENTAL FIGURES

**Supplemental Figure 1.**
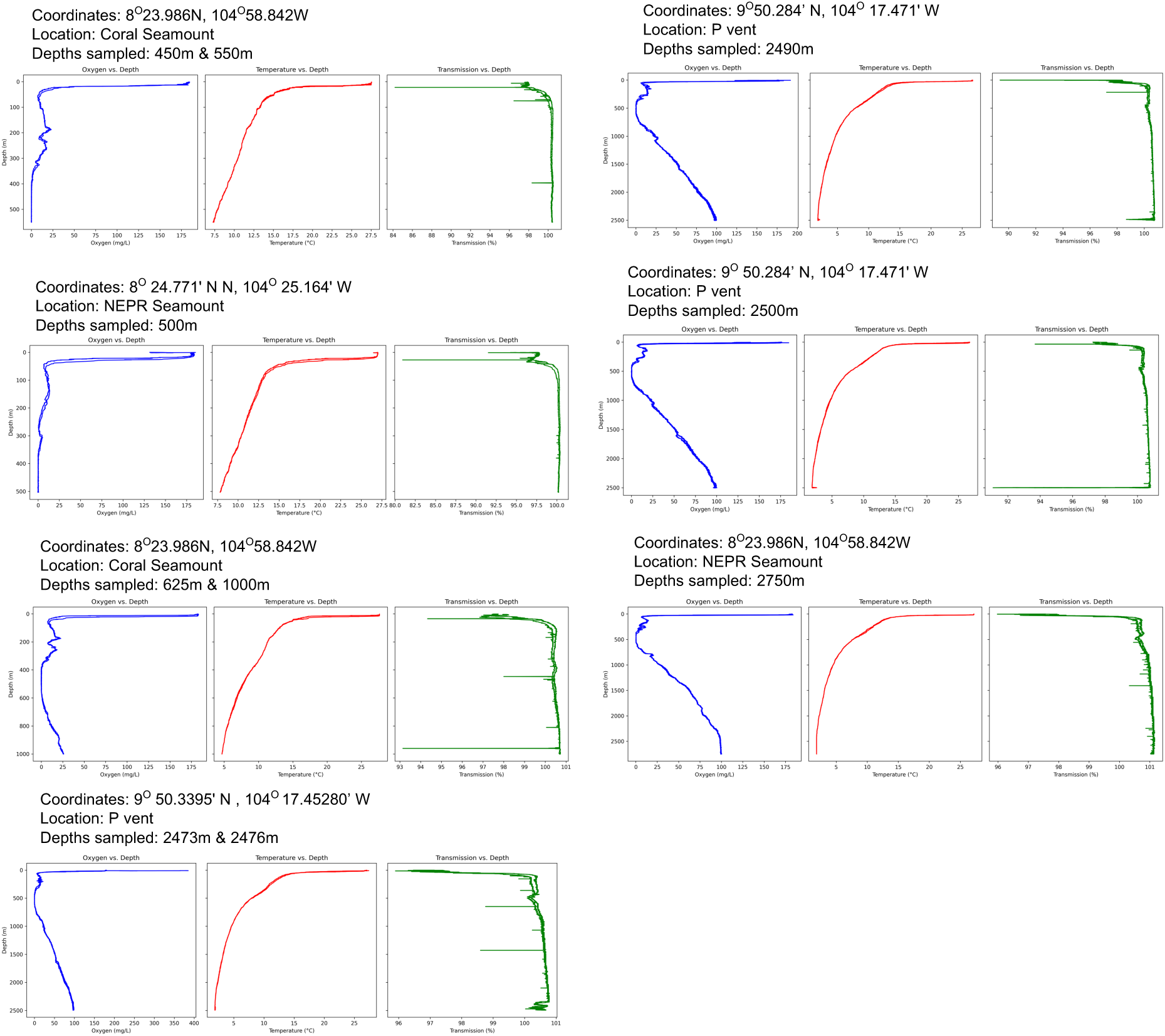
Oxygen, temperature, and transmission profiles of the water column at EPR. One or more depths were sampled on each CTD cast, as indicated on the figure. Also included are the site name and coordinates of the cast.

**Supplemental Figure 2.**
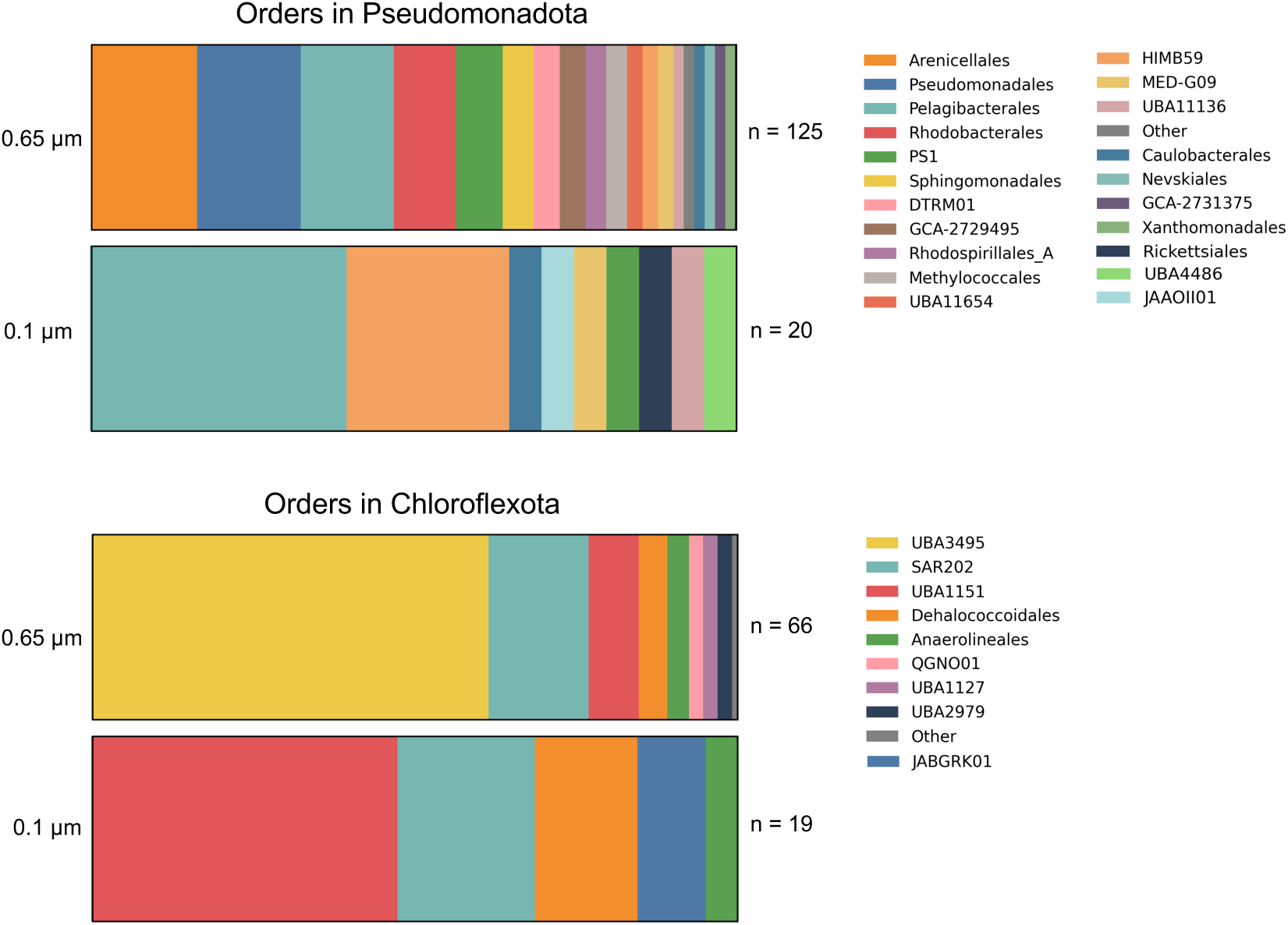
Order-level taxonomic classification of MAGs from phyla Pseudomonadota and Chloroflexota across filter sizes. The number of recovered MAGs in each category is indicated to the right of each bar chart. Orders that consituted <1% of total MAGs were grouped into an “other” category.

**Supplemental Figure 3.**
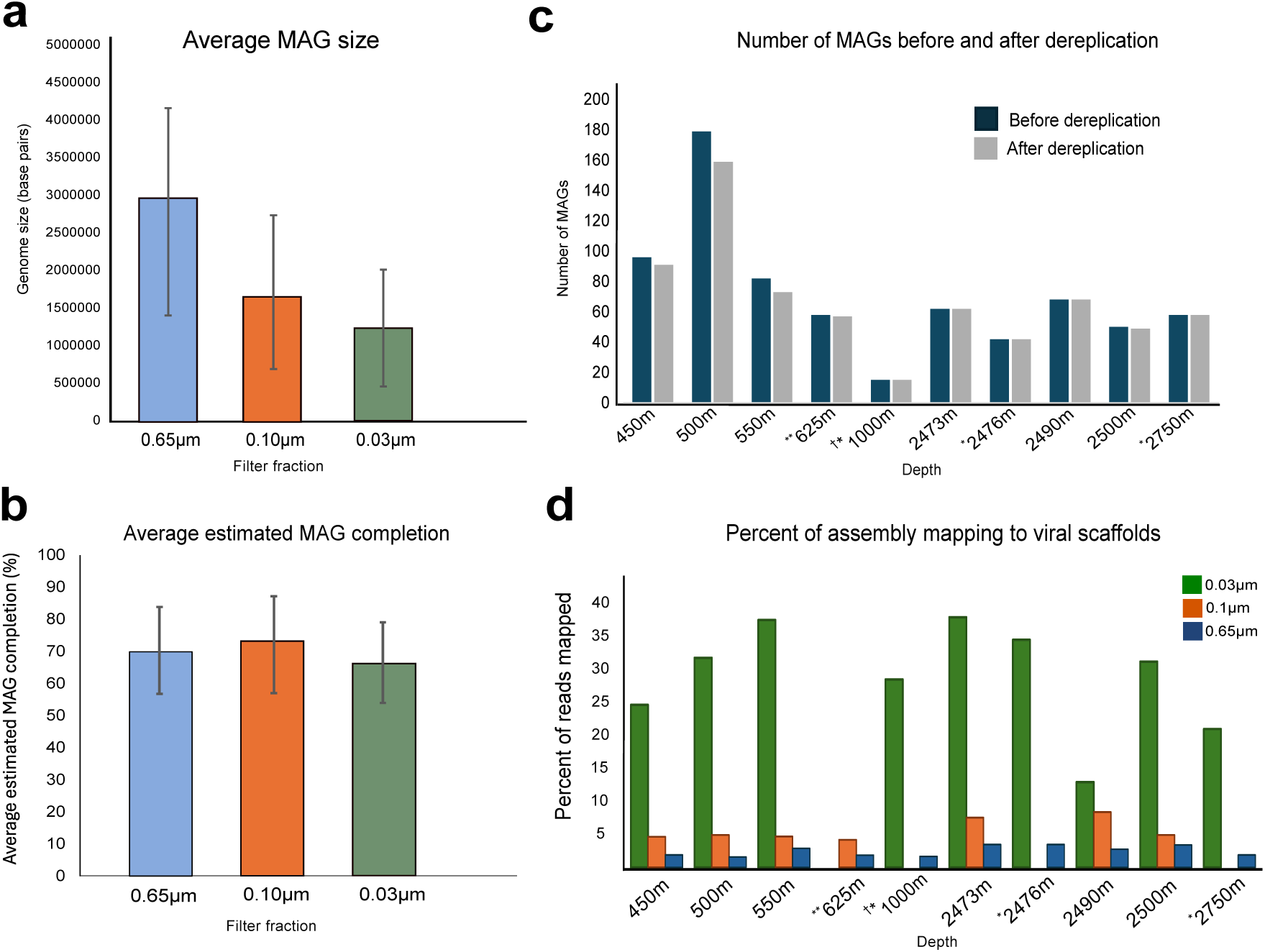
MAG and assembly statistics for EPR metagenomes. a) The estimated size of medium-to-high quality MAGs recovered from each filter size. All sampling depths were considered. Error bars represent standard deviation. b) Average estimated completion of medium-to-high quality MAGs from each filter size as determined by CheckM with all depths considered. Error bars represent standard deviation. c) The number of medium to high-quality MAGs recovered at each depth, considering all filter sizes, before and after dereplication at 95% ANI. On the x-axis, * indicates a 0.1µm assembly was not available for that depth, ** indicates a 0.03µm assembly was not available for that depth, and † indicates the 0.03µm filtered sample was a combination of two depths (625m and 1000m). d) The percent of each assembly that mapped to viral scaffolds identified from the dataset.

**Supplemental Figure 4.**
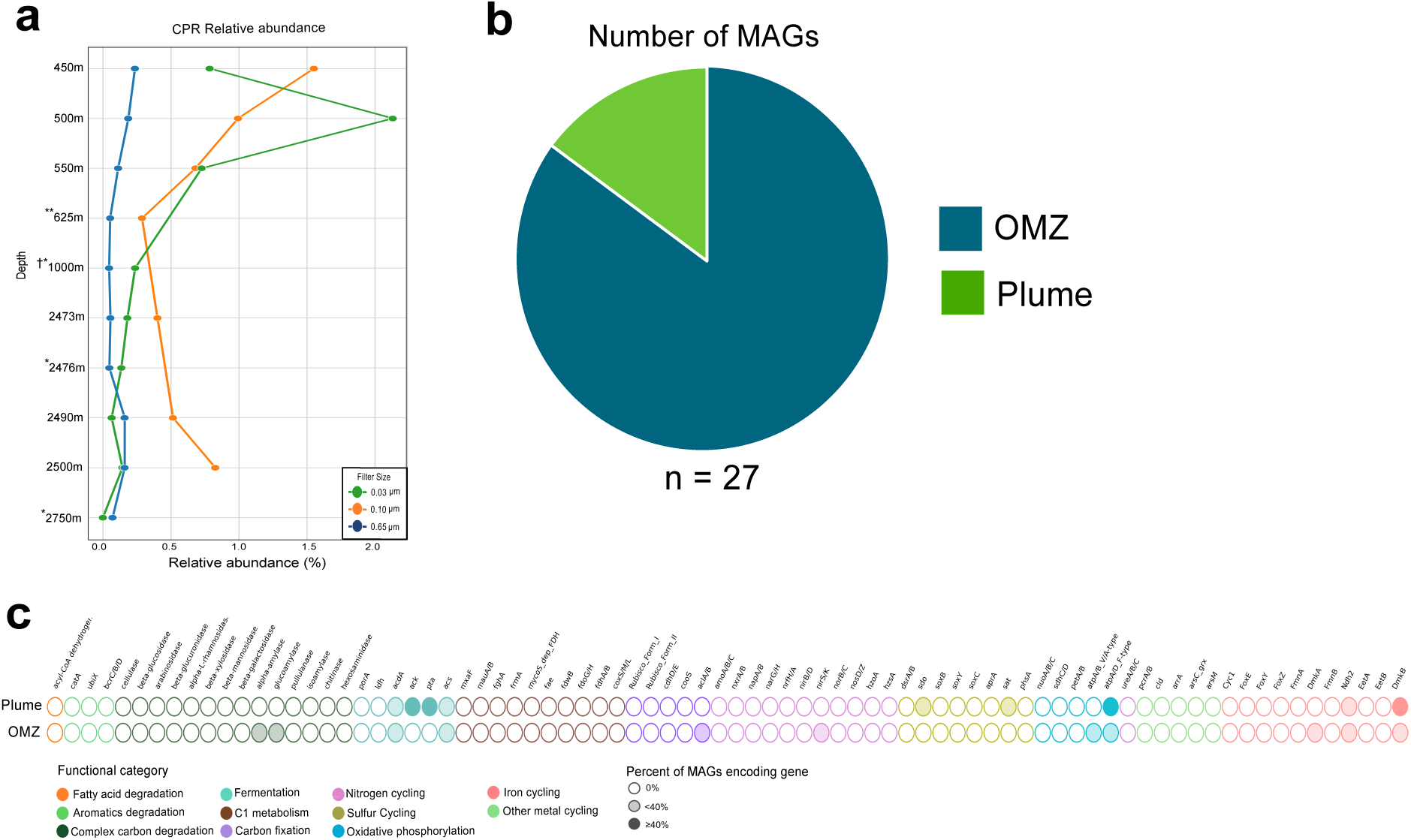
Relative abundance, environmental origin, and metabolism of CPR (Patescibacteria) MAGs. In the relative abundance plot, * indicates a 0.1µm assembly was not available for that depth, ** indicates a 0.03µm assembly was not available for that depth, and † indicates the 0.03µm filtered sample was a combination of two depths (625m and 1000m).

**Supplemental Figure 5.**
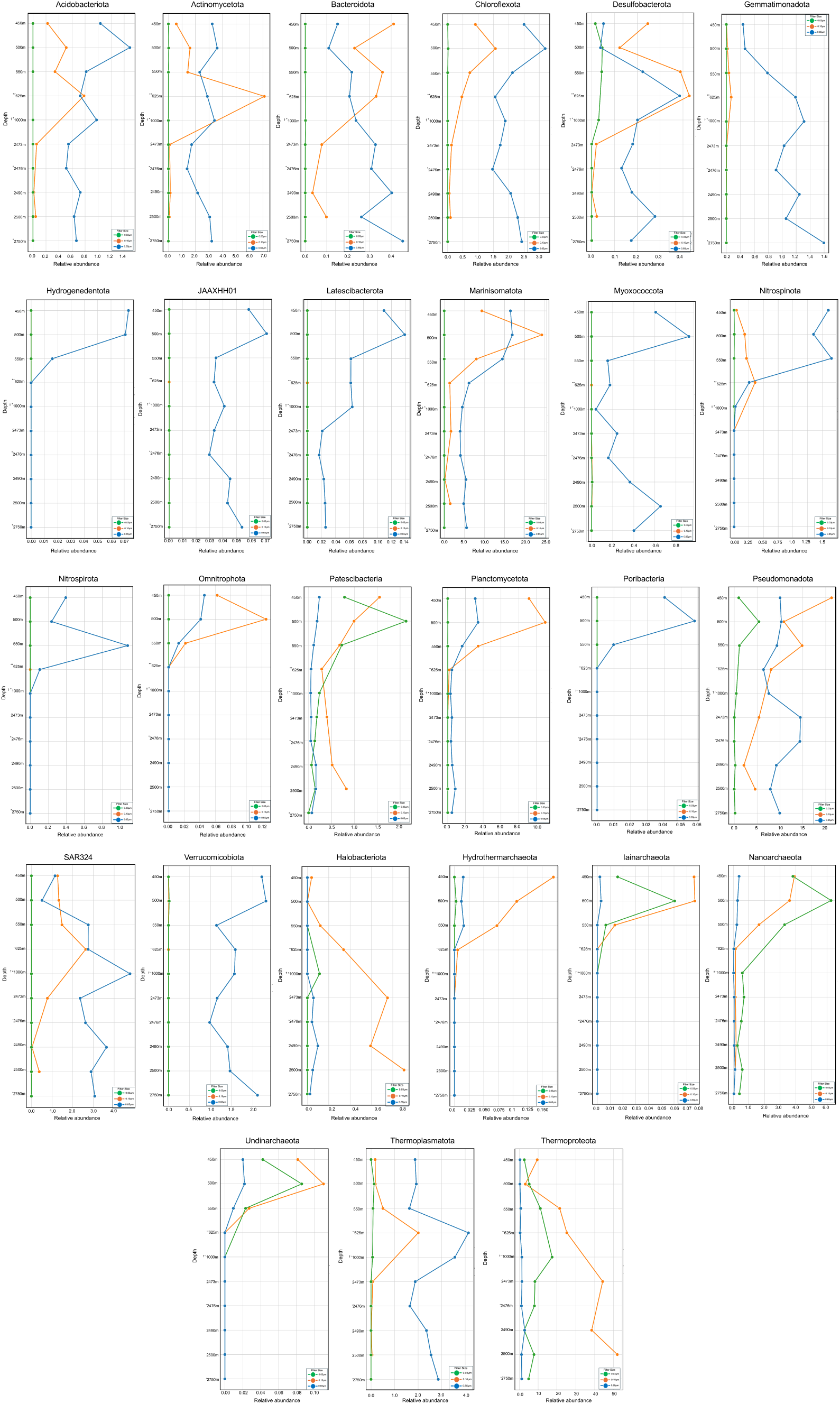
Relative abundance profiles for all bacterial and archaeal phyla in the dataset. Note that the Y-axis scales differ depending on the phylum represented. In the relative abundance plots, * indicates a 0.1μm assembly was not available for that depth and ** indicates a 0.03μm assembly was not available for that depth, † indicates the 0.03μm sample filter was a combination of two depths (625m and 1000m).

**Supplemental Figure 6.**
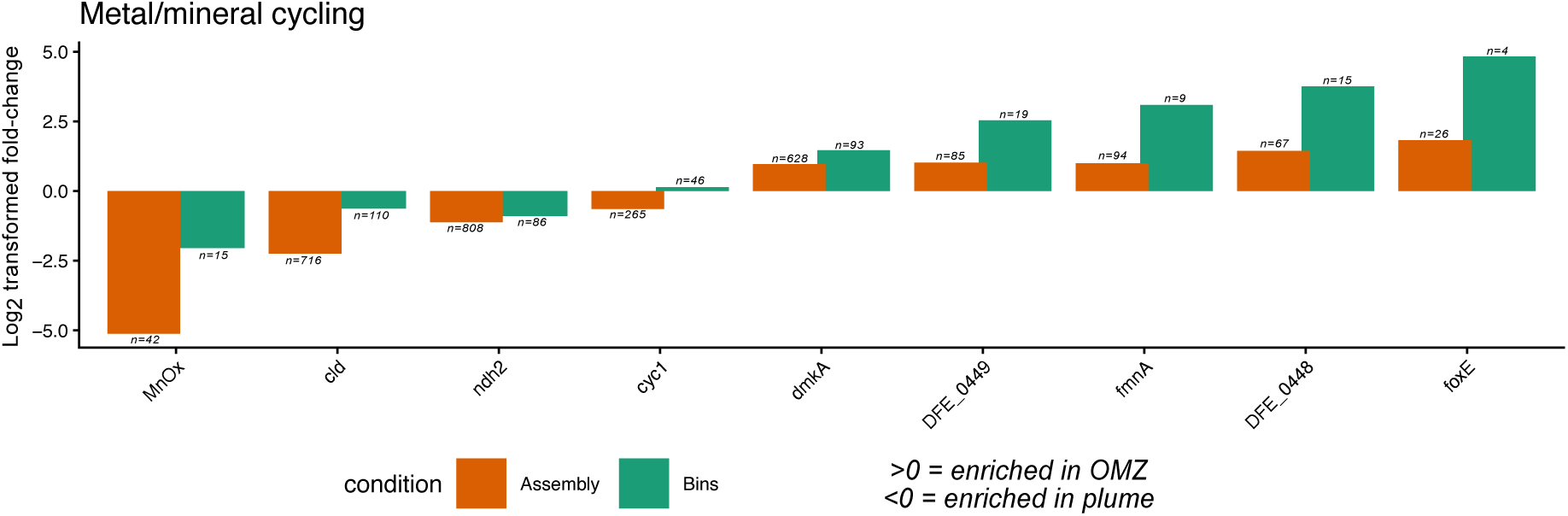
Relative enrichment of genes for metal or mineral metabolism in the plume and OMZ. MnOx = Manganese oxidase; *cld* = chlorite dismutase; *ndh2, dmkA,* DFE_0449, DFE_0448, *fmnA* = iron reduction genes; *foxE, cyc1* = iron oxidation genes

